# Newcastle disease virus induced ferroptosis through p53-SLC7A11-GPX4 axis mediated nutrient deprivation in tumor cells

**DOI:** 10.1101/2021.01.03.424919

**Authors:** Xianjin Kan, Yuncong Yin, Cuiping Song, Lei Tan, Xusheng Qiu, Ying Liao, Weiwei Liu, Songshu Meng, Yingjie Sun, Chan Ding

## Abstract

A number of new cell death processes have been discovered in recent years, including ferroptosis, which is characterized by the accumulation of lipid peroxidation products derived from iron metabolism. The evidence suggests that ferroptosis has a tumor-suppressor function. However, the mechanism by which ferroptosis mediates the response of tumor cells to oncolytic viruses remains poorly understood. Newcastle disease virus can selectively replicate in tumor cells. We show that NDV-induced ferroptosis acts through p53-SLC7A11-GPX4 pathway. The expression of tumor suppressor gene p53 increased after NDV infection, and the expressions of SLC7A11 and SLC3A2 were down-regulated, leading to the inhibition of glutathione synthesis and a decrease in glutathione peroxidase 4 expression. The chemical compound erastin, which induces ferroptosis, also down-regulated glutathione synthase expression and caused lipid peroxide accumulation and cell death. Meanwhile, the levels of intracellular reactive oxygen species and lipid peroxides increased in tumor cells. Ferritinophagy was induced by NDV promotion of ferroptosis through the release of ferrous iron and an enhanced Fenton reaction. Collectively, these observations demonstrated that NDV can kill tumor cells through ferroptosis. Our study provides novel insights into the mechanisms of NDV-induced ferroptosis and highlights the critical role of viruses in treating therapy-resistant cancers.

## Introduction

Ferroptosis, a newly discovered form of regulated cell death, involves iron and reactive oxygen species (ROS), and is characterized by the accumulation of phospholipid peroxides (Chu et al., 2019; Dixon et al., 2012; Doll et al., 2017; Jiang et al., 2015; Yang et al., 2014). The peroxidation of phospholipids and the accumulation of lipid ROS are the key drivers of ferroptosis-induced cell death (Su et al., 2019; Ubellacker et al., 2020). Ferroptosis is initiated through various pathways, including the depletion of intracellular glutathione (GSH) and lipid peroxidation, caused by increases in oxidizing iron and the labile iron pool (Cao and Dixon, 2016; Gao et al., 2019; Hadian and Stockwell, 2020; Muri et al., 2019). GSH is an antioxidant tripeptide consisting of glutamate, cysteine, and glycine (Lv et al., 2019). Recent studies have reported that GSH acts as important regulator of cellular antioxidant defenses. The depletion of cystine and/or GSH results in the iron-dependent accumulation of lethal lipid ROS, which is accompanied by ferroptosis. This process is suppressed by lipophilic antioxidants such as ferrostatin-1 (Dixon and Stockwell, 2014; Hassannia et al., 2019). In contrast, GSH depletion results in the inactivation of GSH-dependent peroxidase 4 (GPX4), a member of the GSH oxidoreductases (GPXs) family, which directly reduces lipid peroxides (Alborzinia et al., 2018). Recent studies have shown that ferroptosis is initiated by the direct depletion or indirect inhibition of coenzyme Q10 via the Squalene synthase mevalonate pathway (Bersuker et al., 2019; Doll et al., 2019; Stockwell et al., 2017).

System X_C_^−^, a cystine/glutamate antiporter, consists of SLC7A11and SLC3A2, and is responsible for maintaining redox homeostasis by importing cystine for the synthesis of the major antioxidant GSH (Xie et al., 2016). Recent investigations have shown that p53 plays a role in the regulation of ferroptosis processes (Jiang et al., 2015; Tarangelo et al., 2018; Wang et al., 2016b; Xie et al., 2017). p53-induced ferroptosis is responsible for tumor suppression through system X_C_^−^, and acetylation is crucial for p53-mediated ferroptosis (Wang et al., 2016b). The function of SLC7A11 is also affected by nuclear factor erythroid 2-related factor 2 (NRF2) and beclin 1 (BECN1). NRF2 upregulates SLC7A11 expression, thereby preventing ferroptosis mediated cell death (Anandhan et al., 2020; Sun et al., 2016; Yu et al., 2020). BECN1 inhibits system X_C_^−^ activity by binding directly to SLC7A11 (Song et al., 2018).

In recent years, there has been renewed interest in ferroptosis in cancer cells (Angeli et al., 2017; Wang et al., 2016a; Yang and Stockwell, 2016). Newcastle disease virus (NDV) is a paramyxovirus that infects large poultry populations, and NDV has been investigated as an oncolytic agent because it replicates selectively in tumor cells (Alexander, 2000; Fiola et al., 2006; Mansour et al., 2011). Many recent studies have shown that NDV promotes oncolytic activity and induces tumor cell death through the caspase-dependent pathway or activates ATM-mediated double-strand break (DSB) signals (Li et al., 2019; Ren et al., 2020). However, there are currently no reports showing whether the oncolytic virus kills tumor cells through ferroptosis. Here, we showed that oncolytic NDV kills cancer cells through ferroptosis via up-regulation of p53 expression and suppression of system X_C_^−^, resulting in the reduction of GPX4 activity. Our study identifies a novel mechanism of oncolytic-virus-induced cell death, and the regulation of ferroptosis may be a promising new way to increase the oncolytic effect of therapies, especially for those therapy-resistant cancers.

## Results

### NDV induces cell death through ferroptosis

Many studies have demonstrated that oncolytic viruses kill cancer cells via various pathways, including apoptosis, autophagy, and pyroptosis (Colunga et al., 2010; Fiola et al., 2006; Koks et al., 2015). However, the role of ferroptosis in oncolytic-virus-induced cell death has not been investigated. Here, we first evaluated whether NDV induces ferroptosis in U251 glioma cells. The ferroptosis inducer, erastin, which inhibits system X_C_^−^ and prevents cystine import, and RSL3, which inhibits GPX4 directly, were used as the positive controls (Dixon et al., 2012). The morphology of cell death caused by NDV was consistent with ferroptosis inducer treatment determined under the microscope (Fig. 1A). Cell death was quantified by assaying lactate dehydrogenase release using a cytotoxic detection kit. Consistently, cell viability decreased significantly upon NDV infection and treatment with the ferroptosis inducer compared with that in control cells (Fig. 1B). The occurrence of free iron in living cells has recently attracted attention because its high reactivity may be related to cell damage or death(Dixon et al., 2012). Ferroptosis initiation depends on the availability of ferrous iron (Fe^2+^)(Hassannia et al., 2019; Masaldan et al., 2018), so we examined whether NDV infection increased the level of intercellular ferrous iron. As expected, NDV increased ferrous iron levels in living cells, which were detected with fluorescent probes (Fig. 1C). Cellular ROS are derived from superoxide radicals, which damage the membranes of cells and organelles (Dixon and Stockwell, 2014; Hassannia et al., 2019). An increased prevalence of ROS was observed in NDV-infected and ferroptosis-inducer-treated cells compared with that in the control groups (Fig. 1D). Accordingly, the oxidative stress marker malondialdehyde (MDA) increased after NDV infection (Fig. 1E). Meanwhile, an increase in lipid peroxidation was confirmed by fluorescence observations using Liperfluo (Green) (Fig. 1F).

**Figure 1.**
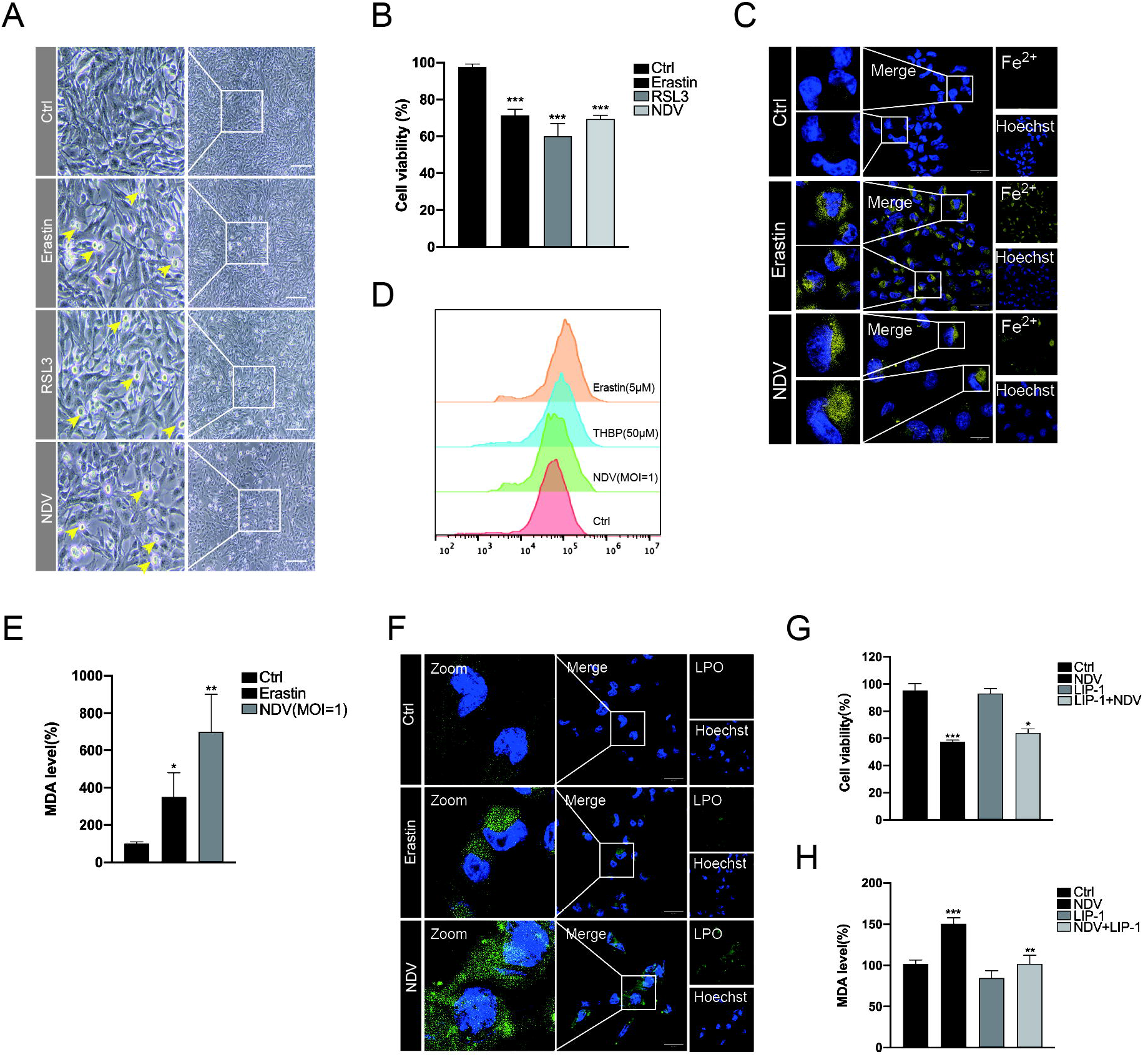
NDV induces cell death through ferroptosis. (A) U251 cells were treated with erastin (5 μM), RSL3 (5 μM), and NDV (MOI = 1) for 24 h, when the images were taken. Scale bars = 1000 &μm. (B) Relative levels of cell viability were assayed by measuring LDH release. (C) Analysis of Fe^2+^ levels in U251 cells after erastin (5 μM) treatment for 24 h using the fluorescent probe FerroOrange (Orange). Scale bars = 20 &μm. (D) Analysis of intracellular ROS levels using DCFDA staining, and flow cytometry of U251 cells treated for 24 h with the compounds indicated. TBHP (50 μM) was used as the positive control. (E) Detection of MDA concentrations in cell lysates according to the kit manufacturer’s instructions (Beyotime, China). (F) Intracellular LPO in U251 cells treated with or without erastin (5 μM) and NDV (MOI = 1) for 24 h were determined with the fluorescent probe FerroOrange (Orange). Scale bars = 20 μm. (G) Cell death was examined with an LDH assay after 24 h of pretreatment with Liproxstain-1. (H) Detection of MDA concentration in cell lysates after 24 h of pretreatment with Liproxstain-1. All experiments were performed at least three times. Significance was analyzed using a two-tailed Student’s t-test. * p < 0.05; ** p < 0.01; *** p < 0.001.

Having shown that oncolytic NDV-induced cell death through ferroptosis, we next tested whether ferroptosis inhibitor blocked U251 cell death in oncolytic NDV-infected cells. Liproxstain-1, a specialized ferroptosis inhibitor, prevents ROS accumulation and cell death in GPX4^−/−^cells (Xie et al., 2016). Our results showed that cell death was noticeably rescued in liproxstain-1-treated NDV-infected cells (Fig. 1G), and that levels of MDA were significantly reduced compared with those in cells only infected with NDV (Fig. 1H). Collectively, these results show that NDV induced cell death through ferroptosis, and the cell death ratio, can be rescued by a ferroptosis inhibitor.

### NDV enables ferroptosis through the suppressor system XC−by activating p53

To determine the mechanism of NDV-induced ferroptosis, we examined whether NDV actives ferroptosis by directly blocking system X_C_^−^. Consistent with previous studies showing that erastin induces ferroptosis by directly targeting system X_C_^−^, our results suggest that NDV suppresses the expression of SLC7A11 and SLC3A2 in a dose- and time-dependent manner (Fig. 2A and B). Activation of p53, a tumor suppressor gene, is known to be required for ferroptosis in cancer cells. Previous studies showed that SLC7A11 protein levels and SLC7A11 mRNA expression are negatively regulated by p53 (Jiang et al., 2015). As expected, our results indicated that NDV also upregulated protein and mRNA levels of p53, accompanied by the downregulation of the protein and mRNA levels of SLC7A11 (Fig. 2A–C). Remarkably, compared with the control group, the reduced GSH levels decreased 5.5- and 8.2-folds in response to erastin treatment and NDV infection, respectively, as measured using a GSSG/GSH quantification kit (Fig. 2D). Accordingly, noticeable time- and dose-dependent reductions in the protein and mRNA levels of GPX4 were observed (Fig. 2A–C). Recent studies have shown that 12-lipoxygenase (ALOX12) is required for p53-mediated tumor suppression through a distinct ferroptosis pathway (Chu et al., 2019). Consistent with the results of Chu et al., ALOX12 increased in a time-dependent manner in NDV-infected U251 cells (Fig. 2E). Taken together, the results presented confirm that the system X_C_^−−^GSH–GPX4 axis is inhibited in response to NDV infection. p53 is probably involved in NDV-induced ferroptosis via the inhibition of the system X_C_^−^.

**Figure 2.**
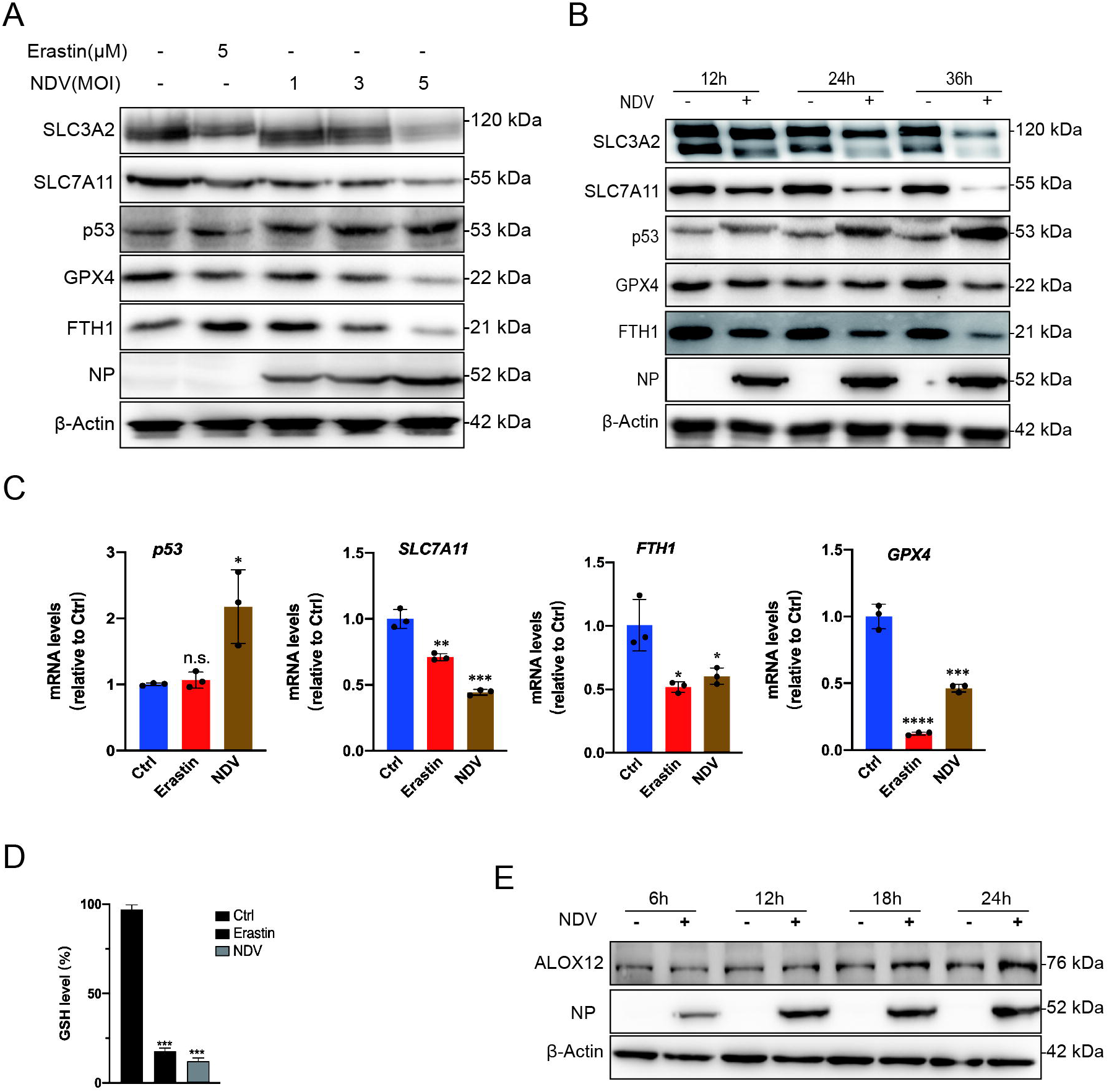
NDV induces ferroptosis by suppressing system X_C_^−^ and activating p53. (A) Cell lysates were analyzed for the levels of p53, SLC7A11, SLC3A2, GPX4, FTH1 and NP, in an MOI dependent manner by western blot. β-Actin was used as the loading control. Erastin (5 μM) was added as a ferroptosis inducer. (B) Western blot analyses of the levels of p53, SLC7A11, SLC3A2, GPX4, NP, and FTH1 in U251 cells, with timepoints indicated. β-actin was served as a loading control. (C) mRNA levels of SLC7A1, p53, FTH1, and GPX4 in U251 cells treated with NDV (MOI = 1). (D) Detection of intracellular GSH concentration after NDV infection. (E) Cell lysates were analyzed for the levels of ALOX12 and NP in a time-dependent manner with western blotting. β-Actin was included as the loading control. All experiments were performed at least three times. Significance was analyzed using a two-tailed Student’s t-test. * p < 0.05; ** p < 0.01; *** p < 0.001.

### p53 plays a positive role in NDV-induced ferroptosis

Given that p53 accumulated significantly after NDV infection, accompanied by the inhibition of the system X_C_^−−^GSH-GPX4 axis, next we investigated the role of p53 in NDV-induced ferroptosis. PFTα HBr (5 μM), a p53 inhibitor, did not affect SLC7A11 and GPX4 protein levels in non-NDV-infected cells, but rescued the protein levels of SLC7A11 and GPX4 in NDV-infected cells (Fig. 3A). We next tested whether PFTα affected MDA production and GSH depletion. Compared with the control group, MDA production was decreased (Fig. 3B) and GSH was rescued after co-treatment with PFTα in NDV-infected cells (Fig. 3C). Moreover, cell death was rescued in PFTα treated NDV-infected cells (Fig. 3D). In accordance with this, lipid peroxide fluorescent intensity was significantly inhibited by PFTα in NDV-infected cells (Fig. 3E). To confirm the intrinsic role of p53 in ferroptosis caused by NDV, small interfering RNA knockdown experiments were performed. As expected, p53 knockdown also rescued the protein levels of SLC7A11 and GPX4 in NDV-infected cells (Fig. 3F). To further illustrate the ways in which p53 plays an important role in NDV-induced ferroptosis, we examined role of p53 in the formation of ROS. As expected, ROS levels were significantly reduced when NDV-infected cells were treated with PFTα (Fig. 3G). Collectively, our inhibitor and knockdown experiments demonstrate that p53 plays a positive role in NDV-induced ferroptosis.

**Figure 3.**
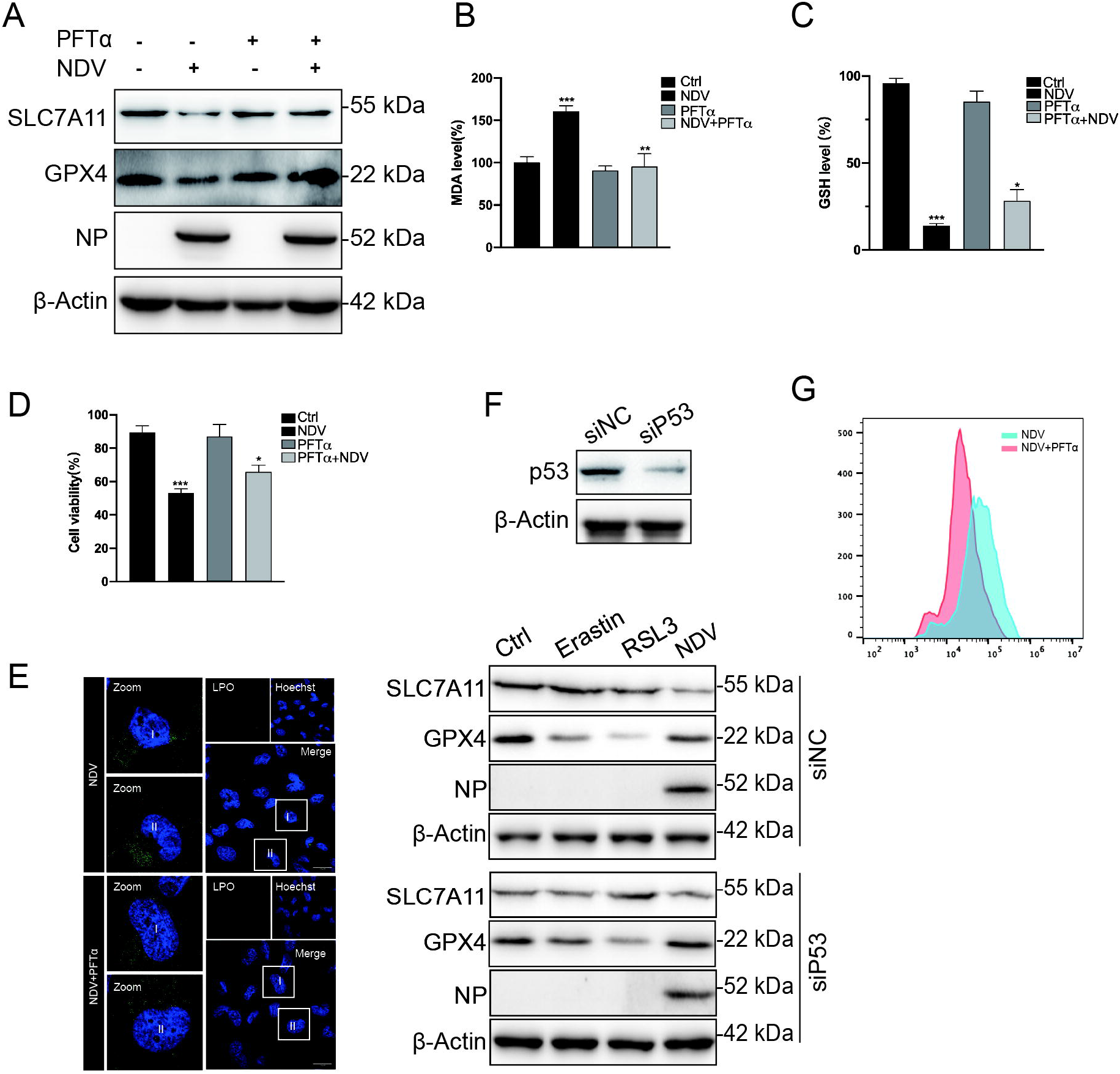
p53 positively regulated ferroptosis caused by NDV. (A) Western blotting analysis of the levels of SLC7A11, GPX4, and NP after PFTα (5 μM) treatment for 24 h in NDV-infected U251 cells. β-Actin was used as the loading control. (B) Relative levels of intercellular MDA were assayed after 24 h of pretreatment with PFTα (5 μM). (C) Detection of intracellular GSH concentrations for 24 h after pretreatment with PFTα. (D) Relative levels of cell viability were assayed by measuring LDH release before treatment with PFTα (5 μM). (E) Intracellular LPO in NDV-infected U251 cells treated with or without PFTα for 24 h was determined with the fluorescent probe Liperfluo (Green). Scale bars = 20 μm. (F) U251 cells with stable knockdown of p53 were treated with erastin (5 μM), RSL3 (5 μM), and NDV (MOI = 5) for 24 h. Western blotting analyses were performed to determine the levels of SLC7A11, GPX4, and NP. (G) Analysis of intracellular ROS levels using DCFDA staining and flow cytometry in U251 cells treated for 24 h with PFTα (5 μM).

### Ferritinophagy induced by NDV contributes to ferroptosis initiation

Ferritinophagy, a new type of autophagy, is characterized by the degradation of ferritin, which promotes ferroptosis through autophagy. Nuclear receptor coactivator 4 (NCOA4), a selective cargo receptor, is involved in the autophagic turnover of ferritin in ferroptosis (Goodall and Thorburn, 2014; Hou et al., 2016). Therefore, we first established whether NDV can induce autophagy in U251 cells. Western blot results showed that NCOA4 levels were significantly reduced in NDV-infected cells, and that viral NP proteins were blocked by the autophagy inhibitor, bafilomycin A (an H-ATPase inhibitor). Next, we determined the LC3-II lipidation levels in NDV-infected cells. LC3 lipidation significantly increased in NDV-infected cells treated with bafilomycin A (20 nM) (Fig. 4A). Excess iron is stored in ferritin, an iron storage protein complex consisting of FTH1 and FTL. Our results showed that FTH1 was shaped and decreased in a dose and time-dependent manner in NDV-infected U251 cells (Fig. 2A–B). Expression of FTH1 mRNA, detected using quantitative polymerase chain reaction (qPCR), was consist with protein levels (Fig. 2D). Previous studies have shown that TFR1, a carrier protein for transferrin in cell membranes, is up-regulated as ferroptosis proceeds. Our investigations showed that TFR1 was not changed significantly in NDV-infected cells.

**Figure 4.**
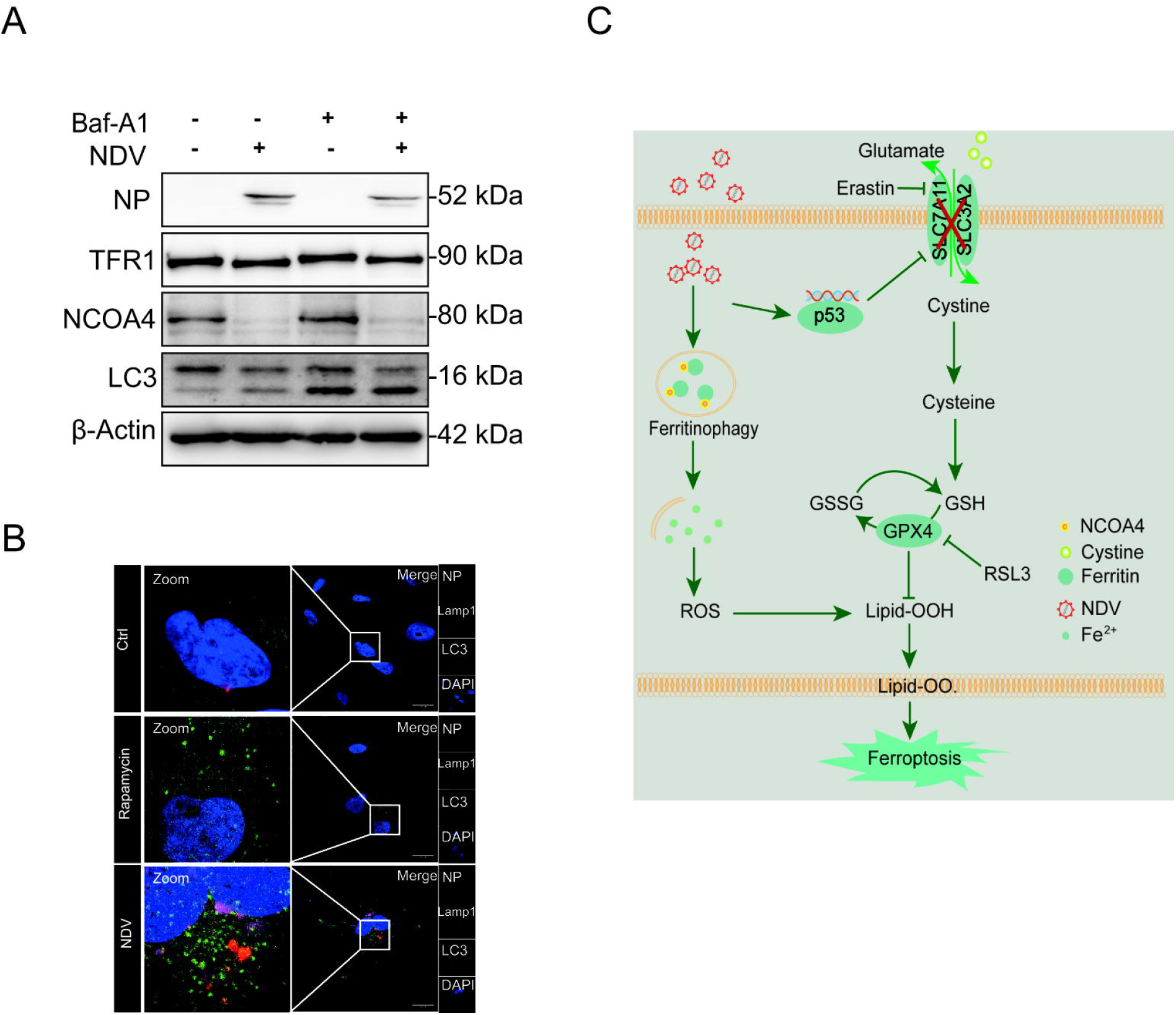
Ferritinophagy induced by NDV contributes to ferroptosis initiation. (A) Western blot analysis of the levels of LC3, NCOA 4, and TFR1 in NDV-infected U251 cells. Baf-A1 (20 nM) was used as an autophagy inhibitor. β-Actin was used as the loading control. (B) Fluorescence microscopy was conducted to assess the ability of NDV to induce the formation of autolysosomes. Staining with Lamp1 (red), and LC3 (green) in U251cells. Scale bars = 20 μm. (A) A model of NDV induces ferroptosis through nutrient deprivation in U251 cells.

Confocal microscopy was used to detect the formation of autophagosomes, seen as green dots. U251 cells were infected with a lentiviral vector encoding an GFP-LC3 reporter. U251 cells stably expressing GFP-LC3 can be used to identify autophagosomes. Our studies showed that NDV significantly increased the number of green dots compared with the control group. These results suggest that NDV accumulates ferrous iron via a decrease in ferritin through autophagy and induces ferroptosis in U251 cells.

## Discussion

Recent studies have shown that regulated cell death is an effective way to kill tumors. Triggering apoptosis cell death using anti-cancer drugs is a classical method of killing cancer cells (Badgley et al., 2020; Jiang et al., 2015; Sun et al., 2016; Tarangelo et al., 2018; Xie et al., 2017; Yang et al., 2014). However, the effects of apoptosis treatment of tumors are limited and the search for new therapies not relying on apoptosis to kill cancer cells will be important for treating therapy-resistant cancers in the future. Ferroptosis, a new form of programmed cell death, is typified by an iron-dependent accumulation of lethal levels of peroxidated lipids, and recent studies have suggested that ferroptosis is a tumor-suppressive mechanism(Stockwell et al., 2017; Yang et al., 2014). Our present study set out to understand the mechanisms of NDV-induced ferroptosis. We provide the evidence that NDV induces ferroptosis through nutrient deprivation in U251 cells, and that activation of p53 was responsible for suppressing system X_C_^−^ and depleting the GSH contribution to lipid peroxidation. Our study suggested that the p53-SLC7A11-GPX4 axis plays a central role in inducing ferroptosis and leading to cancer cell death.

In this study, we showed that NDV suppressed system X_C_^−^, as evidenced by the decrease in SLC7A11 and SLC3A2 protein levels in NDV-infected U251 cells. These results collectively suggest that NDV induces ferroptosis through nutrient deprivation in U251 cells. Consistent with our observations, Dixon et al. showed that System X_C_^−^is a cystine/glutamate antiporter whose activity suppresses ferroptosis in many different cell types (Dixon et al., 2012). Song et al. found that energy stress induces BECN1 activation, and that BECN1 phosphorylation induces ferroptosis by blocking system X_C_^−^ activity via binding to SLC7A11 (Song et al., 2018). In contrast, energy-stress-mediated AMPK activation inhibits ferroptosis via mediated phosphorylation of acetyl-CoA carboxylase and biosynthesis of PUFAs (Lee et al., 2020). Our results show that both SLC7A11 and SLC3A2 were suppressed in NDV-infected cells. Compared with previous studies, our studies suggests that SLC3A2 plays an equally important role in the onset of ferroptosis by suppression of system X_C_^−^function. Recent studies have shown that system X_C_^−^-mediated cystine import is necessary for GSH synthesis. The depletion of cystine, and/or GSH, results in the accumulation of lipid peroxidation products and lethal ROS in an iron-dependent manner (Stockwell et al., 2017).

The activation of p53, a tumor suppressor gene, has been implicated in contributing to the initiation of ferroptosis, and SLC7A11 was identified as a novel p53 target gene. p53 repression of SLC7A11 mRNA expression was observed in the tet-on p53-inducible line (Jiang et al., 2015). In contrast, MDM2 and MDMX induce ferroptosis independently of p53, although MDM2 and MDMX are negative regulators of p53 (Venkatesh et al., 2020). Wang et al. showed that acetylation is crucial for p53-mediated ferroptosis and identified a mouse p53 acetylation at K98 lysine (or at K101 for p53 in humans). The absence of acetylation resulted in the failure of p53 to mediate ferroptosis(Wang et al., 2016b). It has been reported that BRCA1-associated protein 1, another tumor suppressor, induces ferroptosis by repressing SLC7A11 in a similar way to p53 (Zhang et al., 2018). Our present study showed that p53 was upregulated in a dose and time-dependent manner in NDV-infected cells. Inhibition of p53 activation by specific p53 inhibitor, PFTα, blocked NDV-induced ferroptosis. Our results show that MDA production was reduced, and GSH was rescued, after PFTα blocked activation of p53 in NDV-infected cells. SLC7A11 protein levels were also rescued after p53 was suppressed via siRNA. These results collectively suggest that p53 activation plays a critical role in inducing ferroptosis in NDV-infected cells.

As previously reported in the literature, TFR1 is a specific ferroptosis marker and the knockdown of TFR1 makes cells more resistant to erastin-induced cell death (Feng et al., 2020; Shen et al., 2018; Zhang et al., 2020; Zhang et al., 2018). However, our study showed that the protein level of TFR1 was not affected in NDV-infected cells, while ferrous iron production and lipid peroxidation were upregulated after NDV infection in U251 cells. All the results shown above are characteristic of ferroptosis. This is an unresolved issue that requires further investigation of the differences between chemical molecular and virus-induced ferroptosis. Ferrous iron accumulation via ferritin decreases in cells induced Fenton mediation and boosted ROS accumulation to cause lipid peroxidation and to enhance ferroptosis(Liu et al., 2020; Zhou et al., 2020). Accumulating evidence suggests that lysosomes can accumulation iron through degradation of FTH1. NCOA4 has been identified as a cargo receptor (known as ferritinophagy) and recruits FTH1 to autophagosomes while iron release induces ferroptosis (Gao et al., 2016; Hou et al., 2016). Our previous studies showed that NDV triggers autophagy and promotes virus replication in U251 glioma cells (Meng et al., 2012). Here, we have shown that NCOA4 and FTH1 are significantly reduced by autophagy in NDV-infected cells. Hou et al. showed that autophagy induces ferroptosis through the degradation of ferritin (Hou et al., 2016). Consistent with their studies, our results suggest that intercellular ferritin levels were decreased by lysosomes through autophagy enhanced ferroptosis in NDV-infected cells.

Recent studies have shown that ferroptotic responses are often dependent on acyl-CoA synthetase long-chain family member 4. ACSL4 catalyzes the incorporation of PUFAs into membrane phospholipids and enhances ferroptosis (Doll et al., 2017). However, Chu et al. showed that ferroptosis can occur in an ACSL4-independent manner, and that ALOX12 is required for p53-mediated tumor suppression through ferroptosis (Chu et al., 2019). Consistent with this, our results show that ALOX12 is upregulated by NDV infection and ACSL4 activation is not necessary for NDV-induced ferroptosis (data not shown). Recent findings report Ampk-mediated beclin-1 phosphorylation induces ferroptosis (Matthew-Onabanjo et al., 2020; Song et al., 2018). However, Lee et al. showed that PUFAs can be synthesized from the basic building block acetyl coenzyme A carboxylase and that AMPK suppresses ferroptosis by inhibiting ACC. These results suggest that NDV-induced ferroptosis is an ACSL4-independent pathway (Lee et al., 2020). Our present study showed that lipid peroxidation accumulation, as tested using fluorescence probes, and cell death can be significantly suppressed by liproxstain-1. Our findings were limited to the clarification of the mechanism of lipid peroxidation increase in ACSL4-independent manner. Further study is needed to better understand the details of this phenomenon.

In summary, our study demonstrated (1): that NDV-induced ferroptosis acts through nutrient deprivation by suppression of system X_C_^−^; (2) that p53 activation is required for ferroptosis initiation; and (3) the ferritinophagy induced by NDV promotes ferroptosis through the release of ferrous iron (Fig. 4C). Our study first reported that NDV-induced ferroptosis acts via nutrient deprivation, and that oncolytic viruses can kill cancer cells through ferroptosis. Our results explain how NDV infection induces ferroptosis and provide novel insights into new treatments for therapy-resistant cancers.

## Materials and methods

### Cell culture and viruses

Human glioma cells U251 were purchased from the Cell Bank of the Shanghai Institute of Biochemistry and Cell Biology, Chinese Academy of Science (Shanghai, China) (http://www.cellbank.org.cn). U251 cells were cultured in Dulbecco’s Modified Eagle Medium (DMEM) supplemented with 10% fetal bovine serum, 100 μg ml ^−1^penicillin, and 100 μg ml ^−1^streptomycin (all from Thermo Fisher Scientific, Waltham, MA, USA). Cells were cultured at 37°C in an incubator with 5% CO2. NDV (strain Herts/33) was obtained from the China Institute of Veterinary Drug Control (Beijing, China). A stable GFP-LC3 cell line was generated by infecting U251 cells with a lentiviral vector encoding an GFP-LC3 reporter, and human glioma U251 cells (1 × 10^6^–2 × 10^6^ cells) were prepared and infected at a multiplicity of infection (MOI) of 10, with GFP-LC3-overexpressing lentiviruses (GeneChem, Shanghai, China).

### Chemicals and reagents

RSL3 (S8155), liproxstatin-1(Lx-1, S7699), Erastin (S7242), pifithrin-α (PFTα), and HBr (S2929) were purchased from Selleck Chemicals LLC (Houston, TX, USA) and used according to their manufacturers’ instructions. Antibody against β-actin was purchased from Santa Cruz Biotechnology (Santa Cruz, CA, USA). Antibody against NDV nucleoprotein (NP) was prepared in our laboratory. Antibodies against p53 (ab26), SLC7A11 (ab175186), GPX4 (ab125066), ALOX12 (ab11506), and NCOA4 (ab86707) were purchased from Abcam (Cambridge, MA, USA). Antibodies against FTH1 (#4393), TFR1 (#13208), and SLC3A2 (#47213), SignalSilence® Control siRNA (Fluorescein Conjugate) (#6201), and SignalSilence® p53 siRNA I (#6231) were purchased from Cell Signaling Technology (Danvers, MA, USA). DCFDA/H2DCFDA – cellular ROS assay kits (ab113851) were purchased from Abcam (Cambridge, MA, USA). GSSG/GSH quantification kits and Liperfluo were purchased from Dojindo Molecular Technologies, Inc. (Shanghai, China).

### Living cell image analysis

U251 cells were seeded into six-well plates and incubated for 24 h before the experiment began. Inducers or inhibitors were infected with NDV and co-incubated for 24 h. Cells were washed with PBS and then stained with Hoechst 33342 for 10 min at 37°C in the dark, and then washed extensively with PBS. U251 cells were incubated with FerroOrange following the manufacturer’s instructions. Finally, lipid peroxide imaging of the living cells was performed using a fluorescence microscope.

### Western blotting

U251 cells were seeded into six-well plates, infected with NDV (MOI 5), and incubated for 12, 24, and 36 h, or incubated for 24 h in either the presence or absence of inhibitors. Cell samples were washed with PBS and lysed in NP-40 lysis buffer (50 mM Tris-HCl, pH 8.0, 5 mm EGTA, 150 mM NaCl, 2 mM sodium vanadate, 5 mM EDTA, 1% NP-40, and 1 mM NaF) with a cocktail of protease inhibitors (Pierce Chemical, Rockford, IL, USA). Cell lysates were prepared and analyzed for the expression of the indicated proteins using their specific antibodies, followed by treatment with horseradish peroxidase-conjugated secondary antibodies IgG using a chemiluminescence reagent solution kit (Share-bio-Biotechnology, Shanghai, China). β-actin was used as a loading control.

### RNA isolation and quantitative real-time PCR (qRT–PCR)

U251 cells were seeded into six-well plates and total RNA was extracted using TRIzol

^®^ Reagent (Invitrogen, USA) according to the manufacturer’s instructions. cDNA was reverse transcribed from total RNA using expand reverse transcriptase (Roche, USA) and oligo-dT primer. For the quantitative RT–PCR analysis, the following primers were used: p53 forward 5′-CGATCCCGGGCAATACAACT-3′; p53 reverse 5′-TGTGCTGGATGCATCTCTCG-3′; GPX4 forward 5′-CAGTGAGGCAAGACCGAAGT-3′; GPX4 reverse 5′-CCGAACTGGTTACACGGGAA-3′; FTH1 forward 5′-CTGCGCCCTTCTGGAAAATG-3′; FTH1 reverse 5′-GCAACCCCAGGATTTCAGGA-3′; SLC7A11 forward 5′-ACAGGGATTGGCTTCGTCAT-3′; SLC7A11 reverse 5′-GGCAGATTGCCAAGATCTCAA-3′; β-actin forward 5′-GTGGATCAGCAAGCAGGAGT-3′; and β-actin reverse 5′-ATCCTGAGTCAAGCGCCAAA-3′. The mRNA levels of specific genes were calculated using β-actin as the reference gene. All assays were performed in three replicates.

### Lipid peroxidation assay and MDA assay

The lipid peroxidation was determined using Liperfluo (Green), and the detection of MDA concentration in cell lysates was performed strictly to the manufacturer’s instructions (Beyotime, China). Liperfluo, a Spy-LHP analog, is used for lipid peroxide detection and emits intense fluorescence due to lipid peroxide-specific oxidation which allows lipid peroxide imaging using a fluorescence microscope to view living cells. For MDA detection, the thiobarbituric acid included in the kit was added to the supernatants of cell homogenate to form a TBA-MDA mixture, which was then examined spectrophotometrically at 535 nm. All assays were performed with three independent replications.

### GSH/GSHG assay

The intracellular levels of reductive GSH were determined using a GSSG/GSH quantification kit, to detect reductive GSH concentrations in cell lysates, strictly according to the manufacturer’s instructions.

### Statistical analysis

Statistical analyses were performed using GraphPad Prism 8 software (Graph Pad Software, Inc., San Diego, CA, USA). The data were expressed as means ± standard deviation (SD) of at least three independent replications. analyzed using an unpaired Student’s t-test. A p value of < 0.05 was considered statistically significant.

## ABBREVIATIONS

NDV: Newcastle disease virus
NP: Nucleocapsid protein
NCOA4: nuclear receptor coactivator 4
FTH1: ferritin heavy chain 1
FTL: ferritin light chain
TFR1: transferrin receptor protein 1
qPCR: quantitative polymerase chain reaction
AMPK: AMP-activated protein kinase
PUFAs: polyunsaturated fatty acids
BAP1: BRCA1-associated protein 1
ACSL4: acyl-CoA synthetase long-chain family member 4
ALOX 12: 12-lipoxygenase
DCFDA: 2′,7′-dichlorofluorescin diacetate
TBHP: tert-butyl hydroperoxide
MDA: malondialdehyde
LDH: lactate dehydrogenase
ROS: reactive oxygen species
LPO: lipid peroxidation
GSH: glutathione
LIP-1: liproxstain-1
PFTα: pifithrin-α
Baf-A1: bafilomycin A.

## ACKNOWLEDGMENTS

This work was funded by National Natural Science Foundation of China (No.32030108). We also thank Prof. Shengqing Yu (Shanghai Veterinary Research Institute, CAAS) for carefully revising the manuscript draft and giving constructive suggestions.

## References

Alborzinia, H., Ignashkova, T.I., Dejure, F.R., Gendarme, M., Theobald, J., Wolfl, S., Lindemann, R.K., and Reiling, J.H. (2018). Golgi stress mediates redox imbalance and ferroptosis in human cells. Commun Biol 1, 210.

Alexander, D.J. (2000). Newcastle disease and other avian paramyxoviruses. Rev Sci Tech 19, 443–462.

Anandhan, A., Dodson, M., Schmidlin, C.J., Liu, P., and Zhang, D.D. (2020). Breakdown of an Ironclad Defense System: The Critical Role of NRF2 in Mediating Ferroptosis. Cell Chem Biol 27, 436–447.

Angeli, J.P.F., Shah, R., Pratt, D.A., and Conrad, M. (2017). Ferroptosis Inhibition: Mechanisms and Opportunities. Trends Pharmacol Sci 38, 489–498.

Badgley, M.A., Kremer, D.M., Maurer, H.C., DelGiorno, K.E., Lee, H.J., Purohit, V., Sagalovskiy, I.R., Ma, A., Kapilian, J., Firl, C.E.M., et al. (2020). Cysteine depletion induces pancreatic tumor ferroptosis in mice. Science 368, 85–89.

Bersuker, K., Hendricks, J.M., Li, Z., Magtanong, L., Ford, B., Tang, P.H., Roberts, M.A., Tong, B., Maimone, T.J., Zoncu, R., et al. (2019). The CoQ oxidoreductase FSP1 acts parallel to GPX4 to inhibit ferroptosis. Nature 575, 688–692.

Cao, J.Y., and Dixon, S.J. (2016). Mechanisms of ferroptosis. Cell Mol Life Sci 73, 2195–2209.

Chu, B., Kon, N., Chen, D., Li, T., Liu, T., Jiang, L., Song, S., Tavana, O., and Gu, W. (2019). ALOX12 is required for p53-mediated tumour suppression through a distinct ferroptosis pathway. Nat Cell Biol 21, 579–591.

Colunga, A.G., Laing, J.M., and Aurelian, L. (2010). The HSV-2 mutant DeltaPK induces melanoma oncolysis through nonredundant death programs and associated with autophagy and pyroptosis proteins. Gene Ther 17, 315–327.

Dixon, S.J., Lemberg, K.M., Lamprecht, M.R., Skouta, R., Zaitsev, E.M., Gleason, C.E., Patel, D.N., Bauer, A.J., Cantley, A.M., Yang, W.S., et al. (2012). Ferroptosis: an iron-dependent form of nonapoptotic cell death. Cell 149, 1060–1072.

Dixon, S.J., and Stockwell, B.R. (2014). The role of iron and reactive oxygen species in cell death. Nat Chem Biol 10, 9–17.

Doll, S., Freitas, F.P., Shah, R., Aldrovandi, M., da Silva, M.C., Ingold, I., Goya Grocin, A., Xavier da Silva, T.N., Panzilius, E., Scheel, C.H., et al. (2019). FSP1 is a glutathione-independent ferroptosis suppressor. Nature 575, 693–698.

Doll, S., Proneth, B., Tyurina, Y.Y., Panzilius, E., Kobayashi, S., Ingold, I., Irmler, M., Beckers, J., Aichler, M., Walch, A., et al. (2017). ACSL4 dictates ferroptosis sensitivity by shaping cellular lipid composition. Nat Chem Biol 13, 91–98.

Feng, H., Schorpp, K., Jin, J., Yozwiak, C.E., Hoffstrom, B.G., Decker, A.M., Rajbhandari, P., Stokes, M.E., Bender, H.G., Csuka, J.M., et al. (2020). Transferrin Receptor Is a Specific Ferroptosis Marker. Cell Rep 30, 3411–3423 e3417.

Fiola, C., Peeters, B., Fournier, P., Arnold, A., Bucur, M., and Schirrmacher, V. (2006). Tumor selective replication of Newcastle disease virus: association with defects of tumor cells in antiviral defence. Int J Cancer 119, 328–338.

Gao, M., Monian, P., Pan, Q., Zhang, W., Xiang, J., and Jiang, X. (2016). Ferroptosis is an autophagic cell death process. Cell Res 26, 1021–1032.

Gao, M., Yi, J., Zhu, J., Minikes, A.M., Monian, P., Thompson, C.B., and Jiang, X. (2019). Role of Mitochondria in Ferroptosis. Mol Cell 73, 354–363 e353.

Goodall, M., and Thorburn, A. (2014). Identifying specific receptors for cargo-mediated autophagy. Cell Res 24, 783–784.

Hadian, K., and Stockwell, B.R. (2020). SnapShot: Ferroptosis. Cell 181, 1188–1188 e118

Hassannia, B., Vandenabeele, P., and Vanden Berghe, T. (2019). Targeting Ferroptosis to Iron Out Cancer. Cancer Cell 35, 830–849.

Hou, W., Xie, Y., Song, X., Sun, X., Lotze, M.T., Zeh, H.J., 3rd, Kang, R., and Tang, D. (2016). Autophagy promotes ferroptosis by degradation of ferritin. Autophagy 12, 1425–1428.

Jiang, L., Kon, N., Li, T., Wang, S.J., Su, T., Hibshoosh, H., Baer, R., and Gu, W. (2015). Ferroptosis as a p53-mediated activity during tumour suppression. Nature 520, 57–62.

Koks, C.A., Garg, A.D., Ehrhardt, M., Riva, M., Vandenberk, L., Boon, L., De Vleeschouwer, S., Agostinis, P., Graf, N., and Van Gool, S.W. (2015). Newcastle disease virotherapy induces long-term survival and tumor-specific immune memory in orthotopic glioma through the induction of immunogenic cell death. Int J Cancer 136, E313–325.

Lee, H., Zandkarimi, F., Zhang, Y., Meena, J.K., Kim, J., Zhuang, L., Tyagi, S., Ma, L., Westbrook, T.F., Steinberg, G.R., et al. (2020). Energy-stress-mediated AMPK activation inhibits ferroptosis. Nat Cell Biol 22, 225–234.

Li, Y., Jiang, W., Niu, Q., Sun, Y., Meng, C., Tan, L., Song, C., Qiu, X., Liao, Y., and Ding, C. (2019). eIF2alpha-CHOP-BCl-2/JNK and IRE1alpha-XBP1/JNK signaling promote apoptosis and inflammation and support the proliferation of Newcastle disease virus. Cell Death Dis 10, 891.

Liu, J., Kuang, F., Kroemer, G., Klionsky, D.J., Kang, R., and Tang, D. (2020). Autophagy-Dependent Ferroptosis: Machinery and Regulation. Cell Chem Biol 27, 420–435.

Lv, H., Zhen, C., Liu, J., Yang, P., Hu, L., and Shang, P. (2019). Unraveling the Potential Role of Glutathione in Multiple Forms of Cell Death in Cancer Therapy. Oxid Med Cell Longev 2019, 3150145.

Mansour, M., Palese, P., and Zamarin, D. (2011). Oncolytic specificity of Newcastle disease virus is mediated by selectivity for apoptosis-resistant cells. J Virol 85, 6015–6023.

Masaldan, S., Clatworthy, S.A.S., Gamell, C., Meggyesy, P.M., Rigopoulos, A.T., Haupt, S., Haupt, Y., Denoyer, D., Adlard, P.A., Bush, A.I., et al. (2018). Iron accumulation in senescent cells is coupled with impaired ferritinophagy and inhibition of ferroptosis. Redox Biol 14, 100–115.

Matthew-Onabanjo, A.N., Janusis, J., Mercado-Matos, J., Carlisle, A.E., Kim, D., Levine, F., Cruz-Gordillo, P., Richards, R., Lee, M.J., and Shaw, L.M. (2020). Beclin 1 Promotes Endosome Recruitment of Hepatocyte Growth Factor Tyrosine Kinase Substrate to Suppress Tumor Proliferation. Cancer Res 80, 249–262.

Meng, C., Zhou, Z., Jiang, K., Yu, S., Jia, L., Wu, Y., Liu, Y., Meng, S., and Ding, C. (2012). Newcastle disease virus triggers autophagy in U251 glioma cells to enhance virus replication. Arch Virol 157, 1011–1018.

Muri, J., Thut, H., Bornkamm, G.W., and Kopf, M. (2019). B1 and Marginal Zone B Cells but Not Follicular B2 Cells Require Gpx4 to Prevent Lipid Peroxidation and Ferroptosis. Cell Rep 29, 2731–2744 e2734.

Ren, S., Ur Rehman, Z., Gao, B., Yang, Z., Zhou, J., Meng, C., Song, C., Nair, V., Sun, Y., and Ding, C. (2020). ATM-mediated DNA double-strand break response facilitated oncolytic Newcastle disease virus replication and promoted syncytium formation in tumor cells. PLoS Pathog 16, e1008514.

Shen, Y., Li, X., Dong, D., Zhang, B., Xue, Y., and Shang, P. (2018). Transferrin receptor 1 in cancer: a new sight for cancer therapy. Am J Cancer Res 8, 916–931.

Song, X., Zhu, S., Chen, P., Hou, W., Wen, Q., Liu, J., Xie, Y., Liu, J., Klionsky, D.J., Kroemer, G., et al. (2018). AMPK-Mediated BECN1 Phosphorylation Promotes Ferroptosis by Directly Blocking System Xc(-) Activity. Curr Biol 28, 2388–2399 e2385.

Stockwell, B.R., Friedmann Angeli, J.P., Bayir, H., Bush, A.I., Conrad, M., Dixon, S.J., Fulda, S., Gascon, S., Hatzios, S.K., Kagan, V.E., et al. (2017). Ferroptosis: A Regulated Cell Death Nexus Linking Metabolism, Redox Biology, and Disease. Cell 171, 273–285.

Su, L.J., Zhang, J.H., Gomez, H., Murugan, R., Hong, X., Xu, D., Jiang, F., and Peng, Z.Y. (2019). Reactive Oxygen Species-Induced Lipid Peroxidation in Apoptosis, Autophagy, and Ferroptosis. Oxid Med Cell Longev 2019, 5080843.

Sun, X., Ou, Z., Chen, R., Niu, X., Chen, D., Kang, R., and Tang, D. (2016). Activation of the p62-Keap1-NRF2 pathway protects against ferroptosis in hepatocellular carcinoma cells. Hepatology 63, 173–184.

Tarangelo, A., Magtanong, L., Bieging-Rolett, K.T., Li, Y., Ye, J., Attardi, L.D., and Dixon, S.J. (2018). p53 Suppresses Metabolic Stress-Induced Ferroptosis in Cancer Cells. Cell Rep 22, 569–575.

Ubellacker, J.M., Tasdogan, A., Ramesh, V., Shen, B., Mitchell, E.C., Martin-Sandoval, M.S., Gu, Z., McCormick, M.L., Durham, A.B., Spitz, D.R., et al. (2020). Lymph protects metastasizing melanoma cells from ferroptosis. Nature 585, 113–118.

Venkatesh, D., O’Brien, N.A., Zandkarimi, F., Tong, D.R., Stokes, M.E., Dunn, D.E., Kengmana, E.S., Aron, A.T., Klein, A.M., Csuka, J.M., et al. (2020). MDM2 and MDMX promote ferroptosis by PPARalpha-mediated lipid remodeling. Genes Dev 34, 526–543.

Wang, B., Zhang, J., Song, F., Tian, M., Shi, B., Jiang, H., Xu, W., Wang, H., Zhou, M., Pan, X., et al. (2016a). EGFR regulates iron homeostasis to promote cancer growth through redistribution of transferrin receptor 1. Cancer Lett 381, 331–340.

Wang, S.J., Li, D., Ou, Y., Jiang, L., Chen, Y., Zhao, Y., and Gu, W. (2016b). Acetylation Is Crucial for p53-Mediated Ferroptosis and Tumor Suppression. Cell Rep 17, 366–373.

Xie, Y., Hou, W., Song, X., Yu, Y., Huang, J., Sun, X., Kang, R., and Tang, D. (2016). Ferroptosis: process and function. Cell Death Differ 23, 369–379.

Xie, Y., Zhu, S., Song, X., Sun, X., Fan, Y., Liu, J., Zhong, M., Yuan, H., Zhang, L., Billiar, T.R., et al. (2017). The Tumor Suppressor p53 Limits Ferroptosis by Blocking DPP4 Activity. Cell Rep 20, 1692–1704.

Yang, W.S., SriRamaratnam, R., Welsch, M.E., Shimada, K., Skouta, R., Viswanathan, V.S., Cheah, J.H., Clemons, P.A., Shamji, A.F., Clish, C.B., et al. (2014). Regulation of ferroptotic cancer cell death by GPX4. Cell 156, 317–331.

Yang, W.S., and Stockwell, B.R. (2016). Ferroptosis: Death by Lipid Peroxidation. Trends Cell Biol 26, 165–176.

Yu, D., Liu, Y., Zhou, Y., Ruiz-Rodado, V., Larion, M., Xu, G., and Yang, C. (2020). Triptolide suppresses IDH1-mutated malignancy via Nrf2-driven glutathione metabolism. Proc Natl Acad Sci U S A 117, 9964–9972.

Zhang, S., Cao, Y., and Yang, Q. (2020). Transferrin receptor 1 levels at the cell surface influence the susceptibility of newborn piglets to PEDV infection. PLoS Pathog 16, e1008682.

Zhang, Y., Shi, J., Liu, X., Feng, L., Gong, Z., Koppula, P., Sirohi, K., Li, X., Wei, Y., Lee, H., et al. (2018). BAP1 links metabolic regulation of ferroptosis to tumour suppression. Nat Cell Biol 20, 1181–1192.

Zhou, B., Liu, J., Kang, R., Klionsky, D.J., Kroemer, G., and Tang, D. (2020). Ferroptosis is a type of autophagy-dependent cell death. Semin Cancer Biol 66, 89–100.

